# Going with the flow: evidence for phenotypic variation in cooperative plasticity during predator inspection in Trinidadian guppies (*Poecilia reticulata*)

**DOI:** 10.1101/2023.11.01.565091

**Authors:** Sylvia Dimitriadou, Rebecca F.B. Padget, Tegen Jack, Safi K. Darden

**Affiliations:** Centre for Research in Animal Behaviour, Department of Psychology, University of Exeter; Environmental Biology, Department of Biosciences, University of Exeter

**Keywords:** Cooperation, plasticity, Trinidadian guppy, *Poecilia reticulata*, agent-based model, predator inspection

## Abstract

Assortative interactions can be key for the evolution and maintenance of cooperation. The propensity for cooperative behaviour to be met with equal cooperativeness can arise from partner conditional responses or fixed cooperative traits, but also a combination of both; something that has yet to be investigated outside of humans. We explored whether individuals differing in trait cooperativeness also differed in plasticity of their conditional response to partner cooperativeness. To identify when selection may favour high or low plasticity as a function of cooperativeness, we also developed an evolutionary simulation model, where individuals’ probability of cooperating was modelled alongside their plasticity. Our empirical results suggest that guppies (*Poecilia reticulata*) bred to make high cooperative investments in the context of predator inspection exhibit greater conditional response plasticity than guppies bred to make lower investments. Our agent-based model found that more cooperative individuals will show greater plasticity in their propensity to cooperate, compared to less cooperative individuals, except when there are no consequences of nobody cooperating. Combined, our findings show that more cooperative individuals might benefit from a greater capacity to adjust behaviour than less cooperative individuals – this could have implications for assortment by cooperative behaviour.

## 1 Introduction

Across animal species, individuals often cooperate with social partners, incurring costs to provide benefits to others (1–3). Cooperative behaviour amongst unrelated individuals poses an evolutionary conundrum, as natural selection is expected to favour selfish, competitive individuals (2,4–6). It has thus been proposed that an important element in the emergence and maintenance of cooperation amongst non-kin is the assortment of cooperative acts and, therefore, the assortment of a population by cooperation (7), as it ensures the reciprocation of cooperative acts within a network of interacting entities (8). Assortment by cooperation can be a result of flexible behaviour according to conditional rules, where cooperation becomes evolutionarily stable if individuals alter their own cooperative behaviour as a function of past experiences (9); such examples include direct (e.g. ‘Tit-for-Tat’) and generalised reciprocity (10,11). Assortment can also be a result of consistent and repeatable individual differences in cooperativeness, which through non-random, structured interactions can lead to the formation of clusters of cooperative individuals within a social network; such individuals will benefit by associating with each other disproportionately more than they interact with defectors (positive assortment) (12). While often studied separately, assortment by conditional cooperation and positive assortment of cooperative phenotypes are not mutually exclusive, and both may be simultaneously operating within a population (13). Importantly, cooperative phenotypes may differ in responsiveness to the cooperativeness of others – i.e. vary in their behavioural plasticity (14). This suggests that individuals differing in trait cooperativeness may respond to the same social stimuli with differing degrees of variation, something which, to our knowledge, has not yet been investigated in the context of cooperation.

Stability and plasticity in cooperativeness can confer different fitness benefits. For instance, consistent cooperative investment is beneficial if individuals interact repeatedly (15), and may reduce conflict as well as increase specialisation and task sharing in cooperative acts (16). This may result in selection for reduced plasticity in cooperators. That is, if well-connected cooperative individuals defect, their cooperative social neighbours can sever their ties with them, resulting in exclusion and a loss of accrued benefits (17). Conversely, defectors gain no benefits from assorting with other defectors (15) and are unable to establish social ties under mutual agreement (18). Individuals with a lower cooperative propensity are, therefore, expected to have weaker social connections and higher social mobility than cooperative individuals (see 17,18). We might thus expect there to be less selective pressure for consistency in individuals with a phenotypically lower propensity to provide a cooperative investment. Alternatively, the cost of cooperating when others do not might select for high investors to be more plastic in their response to what social partners are doing compared to low investors (i.e. those that are phenotypically less cooperative). This would be particularly relevant in fission-fusion societies where group membership is fluid and individuals have less control over who their interaction partners are. Theoretical and empirical work that would allow us to test these predictions is currently lacking.

In this study, we used the Trinidadian guppy (*Poecilia reticulata*) to explore whether individuals that consistently differ in their propensity to make high or low investments in cooperative acts differ in the degree of plasticity with which they respond when experiencing changes in the cooperativeness of others in their social environment. Guppies cooperate in the context of predator inspection: in the presence of a potential predator, a single individual or a small group of guppies will leave the safety of the shoal to approach and assess the level of the threat posed; when inspectors return to the shoal, this information is transmitted to the remainder of the shoal (19–21). Given that all shoal members benefit from the information collected irrespective of whether they inspected themselves (22–24) and inspecting individuals are thought to incur an increased predation risk (23,25), predator inspection is a model often used to study cooperative behaviour (22,23,25,26). Moreover, the cost of inspection can be shared if inspecting fish form a cooperative partnership (22,26), but experimental work has demonstrated that, in groups of inspecting fish, the individual occupying the lead position incurs a disproportionate risk of predation (23). This means that fish can adjust their level of investment in the cooperative act by moderating their inspection position and our previous work has shown that guppies exhibit consistent individual differences in their tendency to lead during predator inspection (27).

Here, we used two lines of wildtype guppies that had undergone phenotypic selection on the basis of their leading behaviour during a cooperative predator inspection. Individuals originating from the high leading (HL) selection line have been shown to consistently occupy the leading position in pairs of inspecting fish, when compared to individuals from the low leading (LL) line. Importantly, we consider this divergence to reflect the level of ‘investment’ individuals are willing to make during predator inspection bouts rather than differences in risk-taking behaviour, as we have demonstrated that there is no relationship between leading behaviour in this context and boldness following an ostensible predation attempt (27). By manipulating the consistency of cooperativeness in the social environment, we explored whether individuals who show consistent differences in their investment decisions, that is, their leading tendency, differ in the extent to which they adjust their behaviour in response to experiences of cooperation (equal investment) or defection (no investment). Furthermore, to identify conditions under which we may expect to find selection for higher or lower plasticity as a function of individual cooperativeness in the inspection context, we developed an agent-based model, where each individual’s probability of investing in predator inspection as well as its plasticity is modelled in order to understand the emergent evolutionary relationship between cooperativeness and plasticity in cooperative behaviour.

## 2 Materials and methods

### 2.1 Study subjects

Fish tested were laboratory-reared descendants of fish caught from a high predation site in the lower reaches of the river Aripo on the island of Trinidad, TT (N 10°. 39.031; W 61°13.404; 37m altitude) and brought into the lab at the University of Exeter, UK, in March 2008. This study used 160 (80 female, 80 male), adult guppies, that were the fourth generation from breeding lines that had undergone phenotypic selection for high and low leading behaviour during predator inspection (27). Fish were housed in 29cm x 19cm x 15cm tanks in mixed-sex family groups (5 females: 3 males) on a flow-through system at the University of Exeter (12h light: 12h dark cycle) at a temperature of 25oC, and were fed with commercial flake and live food (*Artemia* sp.) twice a day. Fish were anaesthetised and tagged with Visible Implant Elastomer (VIE; Northwest Marine Technology, USA) to allow for individual identification (28) a minimum of 72h before behavioural assays. VIE has been shown not to affect social behaviour in this species (28).

All experimental work was carried out under a UK Home Office project licence (30/3308) and personal licence (I002BDF3F). All methods were performed in accordance with the Animals (Scientific Procedures) Act 1986 (ASPA) and the ARRIVE guidelines.

### 2.2 The behavioural assay

Fish were tested over 4 consecutive days, once per day, using a well-established predator inspection paradigm (26,29,30) in custom-made arenas (Fig. 1). The arenas consisted of two separate inspection lanes divided by an opaque, watertight Perspex barrier. At the end of each inspection lane was a predator compartment, separated by clear Perspex that allowed for the transmission of visual, but not olfactory cues, and placed behind removable opaque Perspex barriers. A realistic model of a guppy predator was placed in the predator compartment, suspended at a height of 2 cm from the bottom of the tank, at an angle facing outwards from the inspection lane. To reduce the possibility of habituation to the predator stimulus, we used models of two common guppy predators, pike cichlids (*Crenicichla alta*) and blue Acara (*Andinoacara pulcher*); in total, 8 predator models were used and were alternated in a semi-randomised order to avoid pseudo-replication. At the other end of the inspection lane was a small compartment placed behind a transparent barrier, containing a model plant. The area close to this plant compartment was the refuge area (see Fig. S1). In the refuge area was a removable, transparent acclimation enclosure (5cm x 14cm). All barriers and enclosures were lifted using a remote pulley system to avoid any disturbance. Cooperation was simulated with the use of a mirror placed lengthwise along the inspection lane to simulate a social partner that inspected alongside the focal, while in the defection condition, a short mirror was placed in the refuge area, to simulate a social partner that would not leave the refuge area to inspect with the focal. To avoid any effects of laterality (29), the long and short mirrors were placed on alternating sides, balanced across individuals and treatments. With this assay we could simulate a fish that invested equally to the focal fish during inspection (full length mirror) or a fish that did not invest at all (short mirror).

Focal individuals were placed into the acclimation enclosure in the refuge zone, where they were left to acclimatise for 9 minutes. At the end of this period, the opaque barrier obstructing the view to the predator compartment was lifted, to allow visual access to the predator. After a further minute, the transparent enclosure was lifted, allowing the fish to freely inspect the model predator for 5 minutes. Individuals were returned to their home tanks after the end of the behavioural assay. Experimental tanks were drained and refilled with fresh water between every trial to eliminate potential odour cues. Trials were video recorded from above using Panasonic HC-V160 cameras (Osaka, Japan).

Videos were analysed using Noldus Observer XT (Version 12) (Wageningen, The Netherlands). Cooperative investment was quantified as the total amount of time an individual spent inspecting the predator (swimming in the inspection lane outside of the refuge area) during the test period (31).Inspection lanes were divided into 10 equidistant zones (including the refuge area), and the time spent in each zone was recorded. The average zone that fish occupied during inspection trials was then calculated. Higher levels of cooperative investment were represented as longer time spent inspecting (i.e. outside of the refuge area), reflecting the amount of inspection, and a closer average proximity to the predator during inspection, reflecting the intensity of inspection. The cooperative behaviour of focal fish in trials where mirrors are used as a simulation for cooperation from social partners during predator inspection has been shown to positively correlate with their cooperativeness when paired with live conspecifics in wild fish (8), validating the use of this paradigm and the behavioural measures outlined above. Behavioural scoring was carried out by three independent observers, blind to the selection line and 4-day experimental treatment; inter-rater reliability was assessed using intraclass correlation coefficient analysis (ICC) (ICC= 0.944, F(_2,22)_=51.84, p <0.001).

### 2.3 Study design

The exposure phase (EP) consisted of the first 3 days (trials), in which fish experienced cooperation or defection. On the fourth day (Test Phase - TP), fish either received the same condition they received during the EP (control treatment), or the opposite condition. Fish from the two selection lines (HL and LL) were distributed equally across treatments. Individuals that did not inspect during 2 or more out of the 3 EP trials were removed from further analysis, as they were not successfully exposed to the experimental condition (4 individuals, across experimental treatments) (see Table S1).

### 2.4 Statistical analysis

We used the package brms (32) in R (33) to fit Bayesian generalised linear mixed models to data for the two behaviour metrics used to describe the propensity for cooperative investment: the time that an individual spent inspecting (measured as a proportion of the 300-second trial time); and the average distance they were from the predator during the inspection time (measured as the weighted average zone during inspection; i.e. riskiness of inspection behaviour). Separate models were used for each behaviour. ‘Fixed’ effects predictors included in the model were: leadership line (two-level categorical; high-leadership/low-leadership); social environment (two-level categorical; cooperation/defection); trial day (four-level categorical; days 1, 2, 3, 4); sex (two-level categorical, female/male); predator stimulus (two-level categorical, blue acara/pike cichlid); lane (eight-level categorical variable describing which test lane was used). Predator stimulus and lane were counterbalanced in the experimental design, but we included them in the model to allow more precise effect size estimates. We also included a three-way interaction between the line, social environment, and treatment condition. Treatment was a categorical variable with four levels, which described the sequence of social conditions that the individual experienced over the exposure and test phases (Table 2). The interaction between line, social environment, and treatment term would allow us to determine whether the lines showed different responses when their social environment in the treatment phase did or did not match the one experienced in the exposure phase and whether this was in a positive direction (cooperation following serial defection) or a negative direction (defection following serial cooperation). The treatment was used only in the interaction term because this meant that it described whether individuals experienced the same or different behaviour from social partners than what they might expect from their previous experience; if it were included as a term on its own it would capture aspects of both trial day and social environment, meaning that it would be uninterpretable. ‘Random’ effects terms for the housing tank ID (categorical with 30 levels) and fish ID (categorical with 163 levels) were also included to account for any non-independence within housing tanks, and in individual behaviour, respectively.

**Table 2.**
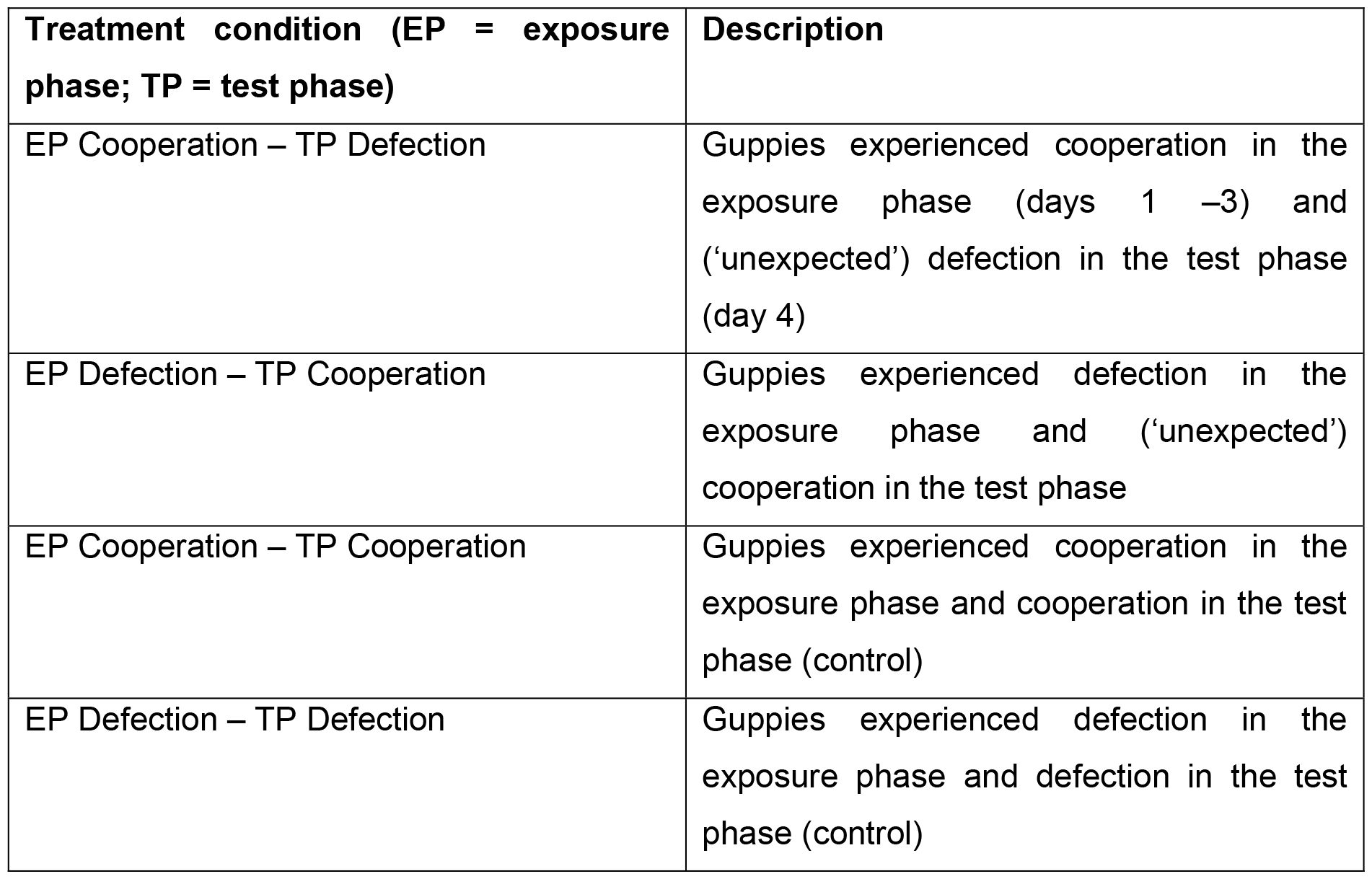
Description of treatments.

For clarity of interpretation, we report marginal effects of parameters (rather than coefficient estimates). We give this as the median of the difference between the posterior draws from each parameter and calculate 50%, 89%, and 95% confidence intervals of these effects using the R package marginaleffects (34). We report these results on the scale of the response variable.

### 2.5 Simulation model

To formalise our verbal model and identify the conditions under which we might expect to have found our experimental results, we developed a simulation model. The model runs by initialising a population of individuals, each of which is given an investment trait and a plasticity trait, which represent the investment an individual makes in inspection and an amount by which they can adjust this investment, respectively. The investment and plasticity are determined for each fish independently by randomly drawing values from a uniform distribution bounded at 0.3 and 0.7 for the investment, and between 0 and 1 for the plasticity. For each ‘step’ of the model run, a focal individual is chosen. It is then assigned a different payoff depending on its investment in cooperation and the investment of other individuals: if the mean investment of other individuals plus noise (drawn from a normal distribution with mu = 0 and sigma = mean of the group’s plasticity) is less than or equal to the focal individual’s investment plus noise (∼ N(0, individual’s plasticity)), the individual receives a payoff, 1 - cost of inspecting (c) * investment / mean investment of the group plus noise; if the mean investment of all individuals is less than a (constant) threshold (0.3), the focal receives a payoff, 1 – cost of lack of investment (a); if the focal makes less than the mean investment, it receives payoff, 1. These payoffs are approximately equivalent to a continuous N-person snowdrift game with cost-sharing. If the payoff received is below a randomly generated number between 0 and 1, then the focal either adjusts its future investment, by its plasticity parameter, or dies – this is determined by chance and there is equal probability of each outcome. When an individual dies, it is immediately replaced by choosing a random parent from the population (not including the focal), and giving the new fish its mean and standard deviation attributes plus some noise drawn from a normal distribution (noise ∼ N(0, 0.1)) to represent genetic mutation.

The model was run 100 times for 50,000 steps for each combination of parameter values for c and a for 1 ≤ c < 2 and 0 ≤ a < 1. When c is high, this represents a high cost of cooperation; when a is high, this represents a high cost incurred if insufficient investment is made by the whole group. To measure the relationship between cooperativeness and plasticity, we calculated the Pearson correlation coefficient between cooperativeness and absolute plasticity from focal fish in the final 100 timesteps of each run to find a population-level relationship between cooperativeness and plasticity.

## 3 Results

### 3.1 Empirical experiment

#### 3.1.1 Time spent inspecting

##### Plasticity

To measure plasticity in the test condition, we compared behaviour on the final trial day to behaviour on the previous days. We found that high-leadership (HL) fish inspected for almost 40 seconds longer (out of 300 seconds) on the final trial day when partners switched from defection to cooperation (DDD/C). There was no evidence of this change in behaviour in the low-leadership (LL) guppies, and there was no evidence of a change in behaviour in either line when partners switched from cooperation to defection (Table S2) (Figure 2).

**Figure 2.**
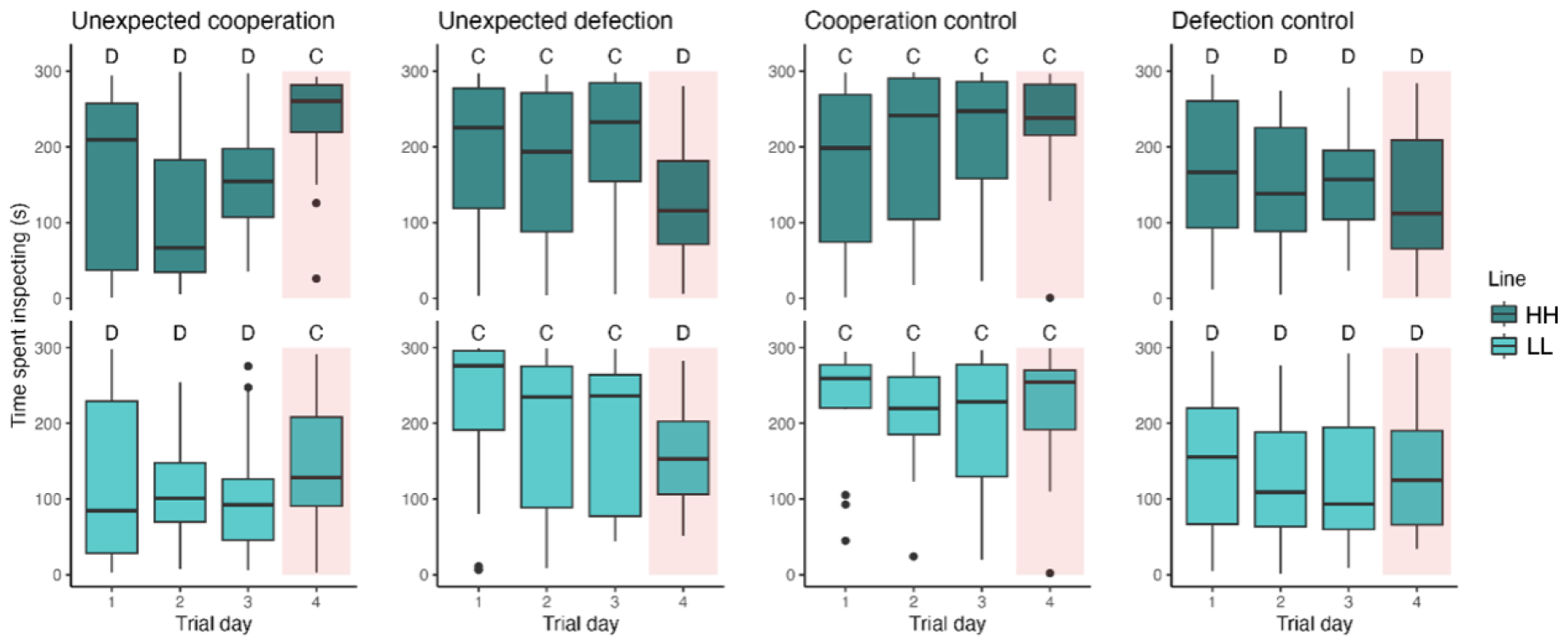
Time spent inspecting: plasticity. The top row shows the high-leadership and the bottom row shows the low-leadership line. The red rectangles highlight the final trial day – the ‘test phase’. The labels above each box indicate the social environment that the group experienced on that day: cooperation (C) or defection (D). Fish in the high-leadership line spent more time inspecting when they experienced partners who switched to cooperation (top left), while fish in the low-leadership line did not (bottom left).

##### Comparison to baseline

To determine whether guppies responded to their partner switching behaviour from previous trial days compared to how they would respond to this behaviour if it had been experienced previously, we compared behaviour on the final trial day in the treatment condition to behaviour in the control condition (in which fish experienced the same behaviour from their partners on all days). There was some evidence that HL guppies invested more in inspection when cooperation was unexpected (i.e. exceeded the baseline investment when partners switched from defection to cooperation) than they did when experiencing continuous cooperation from a partner. In the LL guppies, there was evidence that individuals invested less under unexpected cooperation (i.e. did not reach the baseline investment when partners switched from defection to cooperation) than when cooperation was continuous. There was evidence that HL guppies invested less in inspection when defection was unexpected (i.e. did not reach the baseline investment when partners switched from cooperation to defection) compared to when experiencing continuous defection; for LL guppies, there was also a slight negative marginal effect, but credible intervals overlapped zero at the 50% level (Table S2) (Figure 3).

**Figure 3.**
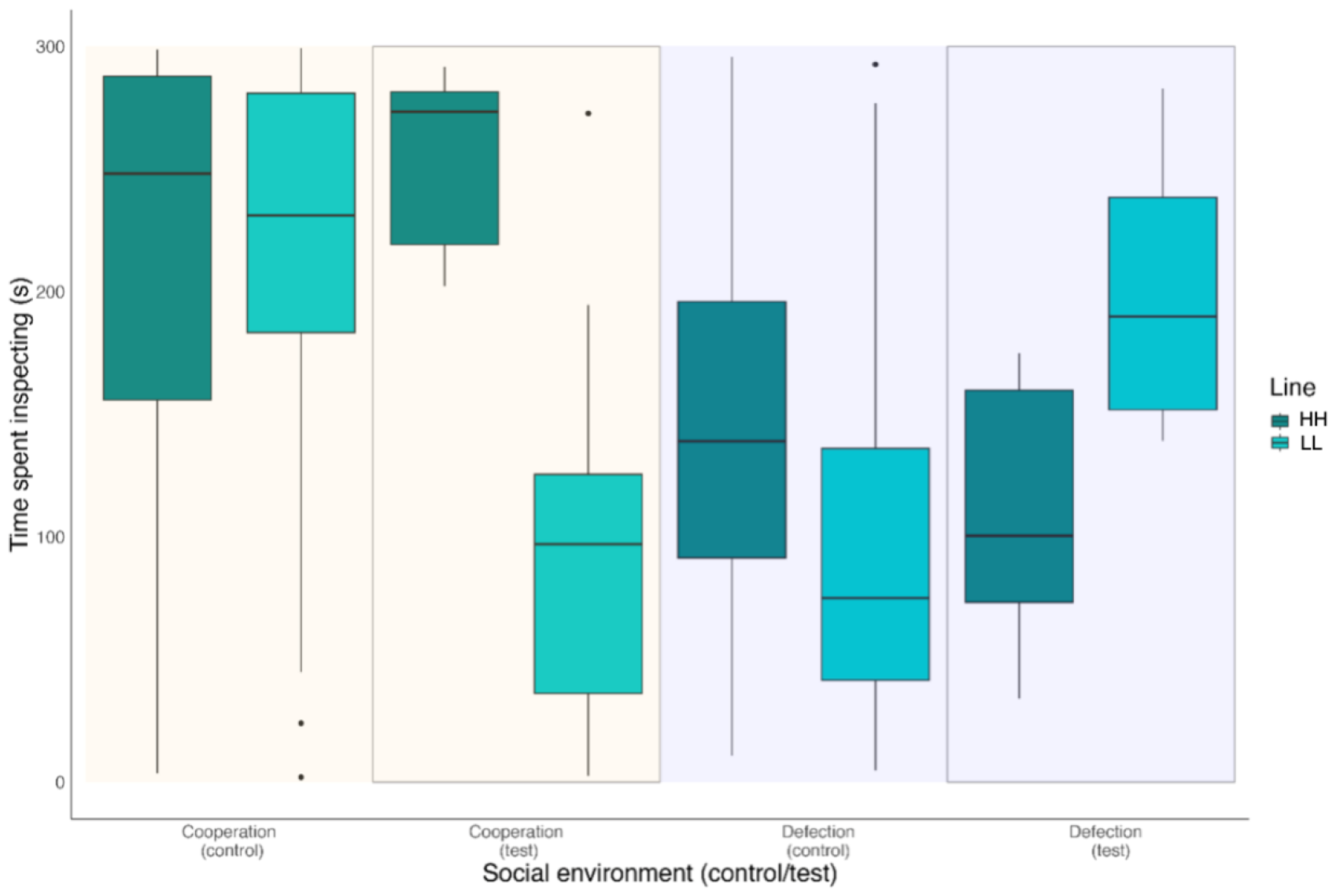
Time spent inspecting: comparison to baseline. Boxes show the time (in seconds) that guppies spent inspecting. Line is shown by the colours of boxes – dark green shows high-leadership lines; light green shows low-leadership lines. Boxes on an orange background show trials in which the test fish ostensibly perceived cooperation; boxes on a blue background show trials in which the test fish ostensibly perceived defection. The grey outlines highlight the test trials in which the guppies experienced a switch in partner behaviour relative to the previous three inspection events. This figure shows that the lines responded differently when partners switched to cooperation, but not when they switched to defection.

Results from other model parameters are given in Table S4 in the Supplemental Information.

#### 3.1.2 Distance to predator stimulus

##### Plasticity

There was no evidence that either HL or LL guppies changed the distance at which they inspected over consecutive trial days in response to their partner switching to either cooperation or defection (Table S3) (Figure 4).

**Figure 4.**
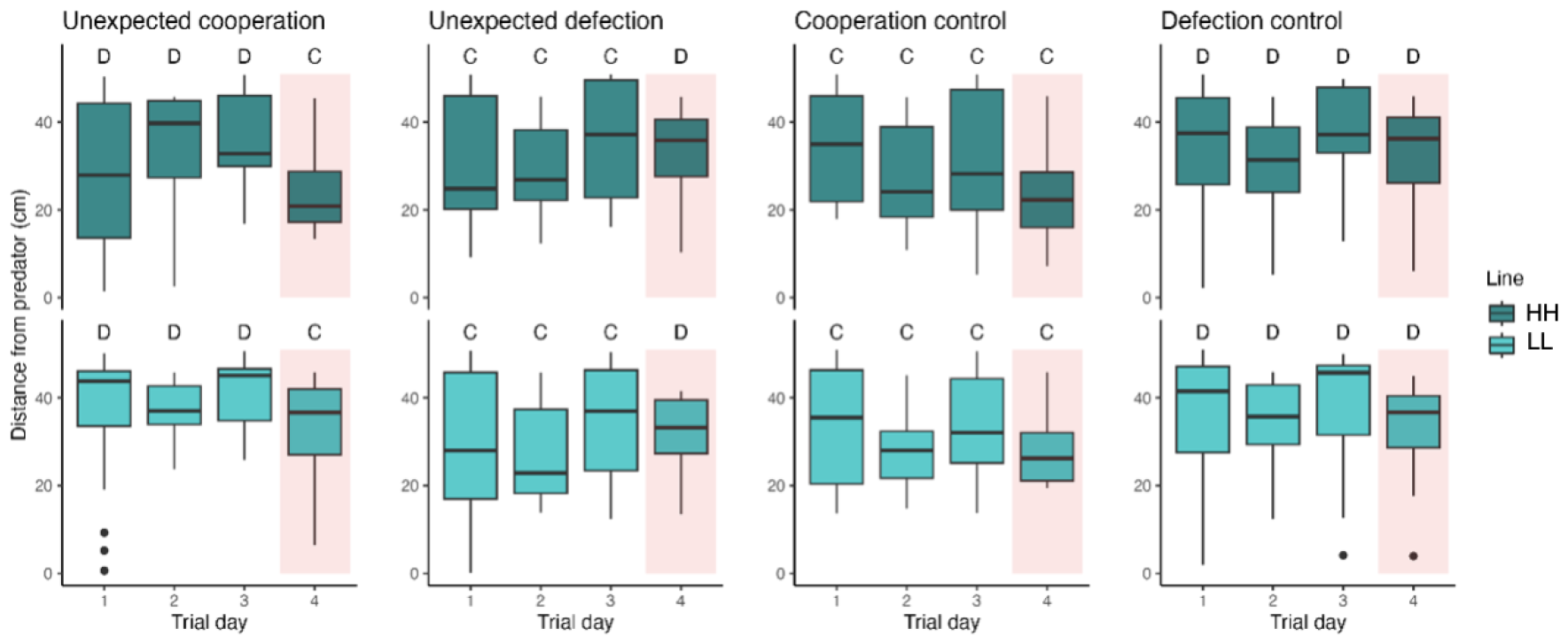
Distance from predator: plasticity. The top row shows the high-leadership line and the bottom row shows the low-leadership line. The red boxes highlight the final trial day – the ‘test phase’. The labels above each box indicate the social environment that the group experienced on that day: cooperation (C) or defection (D). Lines did not change how close they inspected in response to changes in conspecific behaviour.

##### Comparison to baseline

When guppies experienced unexpected cooperation, HL guppies inspected closer than they did when experiencing continuous cooperation (i.e. exceeded the baseline investment when partners switched from defection to cooperation). The opposite was true for LL guppies, who inspected from further away when cooperation was unexpected compared to continuous (i.e. did not reach the baseline investment when partners switched from defection to cooperation). Both lines appeared to inspect from further away when defection was unexpected compared to when they experienced continuous defection (i.e. did not reach the baseline investment when partners switched from cooperation to defection) (Table S3) (Figure 5).

**Figure 5.**
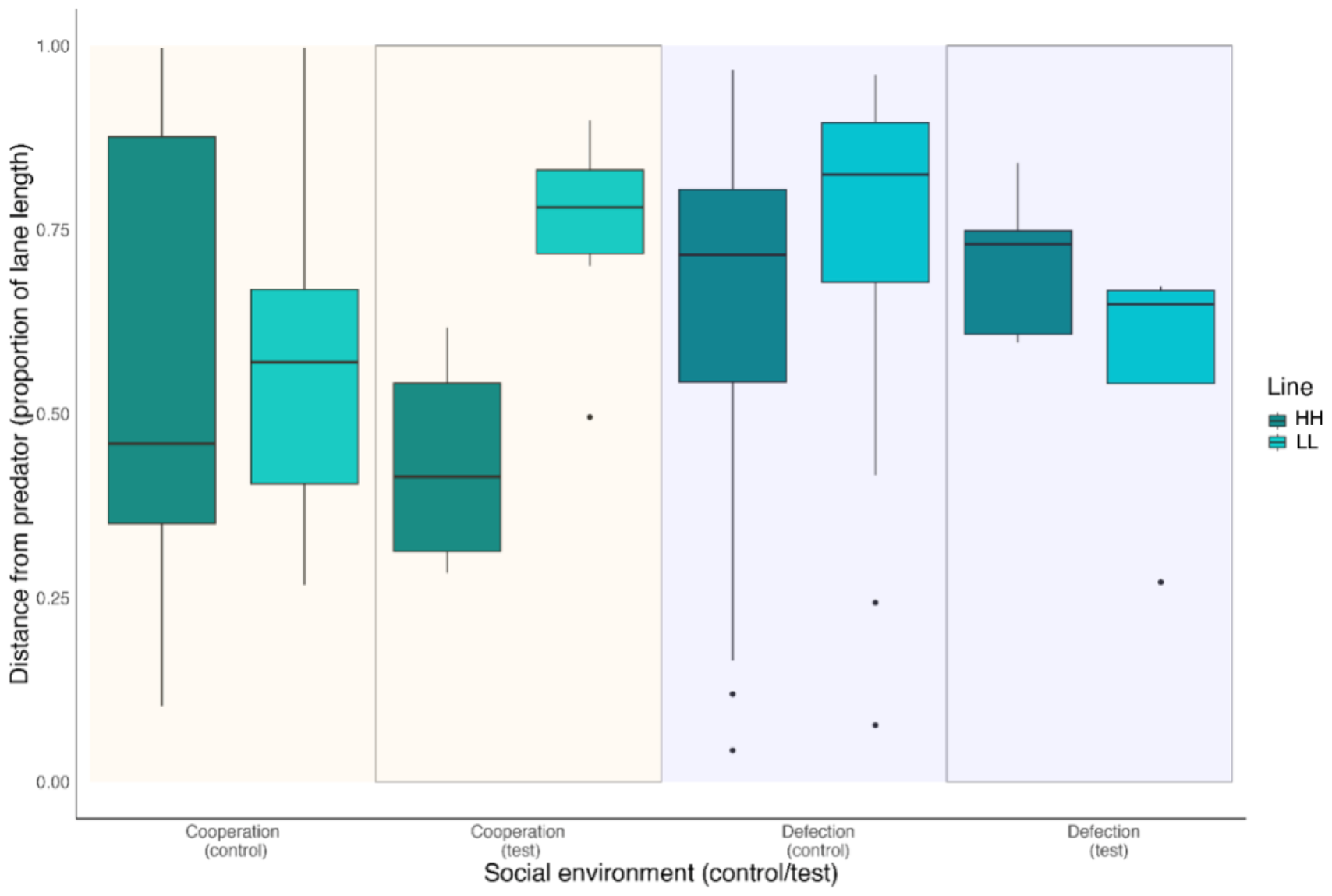
Distance from predator: comparison to baseline. Boxes show the distance as a function of lane length that guppies maintained to the predator (1 = furthest possible distance, 0 = closest possible distance). Line is shown by the colours of boxes – dark green shows high-leadership lines; light green shows low-leadership lines. Boxes on an orange background show trials in which the test fish ostensibly perceived cooperation; boxes on a blue background show trials in which the test fish ostensibly perceived defection. The grey outlines highlight the test trials in which the guppies experienced a condition where social partners switched behaviours on the trial day. This figure shows that the lines perhaps responded differently to a switch to cooperation (orange background, in rectangle), but did not differ in their response to a switch to defection.

#### 3.1.3 Exposure phase comparisons

During the exposure phase, high- and low-leadership guppies did not differ in their inspection time or distance. This was true regardless of whether they experienced a cooperation or defection stimulus during this phase. While high-leadership guppies appeared to respond similarly to cooperation and defection in the exposure phase, there was some weak evidence that the low-leadership guppies invested more in inspection when they experienced cooperation than they did when they experienced defection (marginal effect = +33 seconds; 50% HDI = 4.1, 19; 89% HDI = -5.5, 28).

### 3.2 Simulation model result

We found that generally there was a positive correlation between level of cooperative investment and plasticity, except when there was no cost of non-inspection (a = 0), when there was a negative relationship. This indicates that more cooperative individuals generally evolved to have slightly greater plasticity (Figure 6). This suggests that more cooperative individuals should adjust their cooperative investment more when there are consequences (i.e. risks) to non-cooperation.

**Figure 6.**
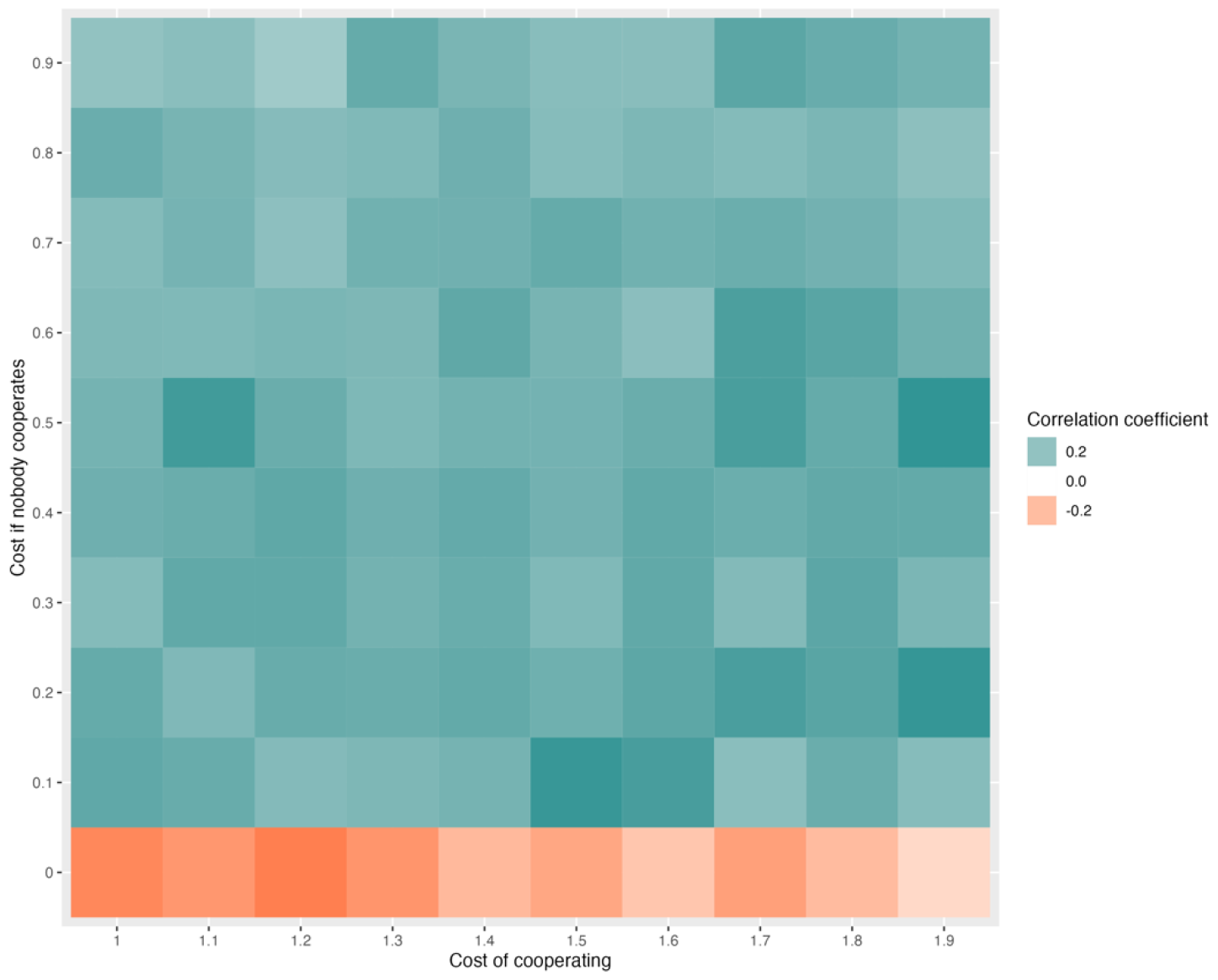
Heatmap of correlation coefficients as the cost multiplier of cooperating and the cost if nobody cooperates change. Typically, a positive correlation between cooperation and plasticity emerges, meaning that, on a population level, greater cooperativeness is accompanied by greater plasticity. However, there is a negative relationship when there is no cost to cooperation. Colour represents the correlation between investment and plasticity of focal individuals in the final 100 timesteps (of 50,000); correlation coefficients shown are the means of 100 independent runs of the model.

## 4 Discussion

In this study, we tested whether guppies bred to make higher or lower investments during cooperative predator inspection differed in their responsiveness to changing cooperativeness in a simulated social partner. We found that more cooperative guppies showed greater evidence of plastically adjusting their cooperative investment depending on the behaviour of their social partner than less cooperative guppies. More cooperative guppies invested more when their partner switched from defecting to cooperating; less cooperative guppies showed no evidence of adjusting their behaviour in consecutive trials. As further evidence of their plasticity, more cooperative guppies actually appeared to exaggerate their behaviour when their partner switched from defection to cooperation, inspecting closer and for longer than the control group, which experienced cooperation on all trial days. In contrast, less cooperative guppies inspected for less time when experiencing unexpected cooperation than did the control group, which experienced continuous cooperation, and thus did not adjust to the change. There was some evidence that less cooperative guppies inspected from further away when experiencing unexpected behaviour from a partner, regardless of whether their partner switched to cooperation or defection. The results of our agent-based model support our empirical results, finding that a positive correlation can emerge between cooperativeness and plasticity (given that there is some cost when nobody cooperates). The simplicity of our model highlights that this correlation can emerge even under very simple behavioural rules.

Our empirical findings suggest that more cooperative individuals show greater plasticity than less cooperative individuals. Given that individuals inspecting alone assume higher risk than those joined by social partners (e.g. 23), individuals more inclined to cooperate might benefit more from adjusting to a partner’s behaviour, compared to less cooperative individuals. The high flexibility of cooperating individuals might enable them to avoid exploitation from defectors, whilst maintaining a high propensity for cooperating when social conditions are favourable (as in a tit-for-tat strategy). Under these dynamics, assortment could emerge through the conditional reciprocation of cooperation (9), leading to repeated interactions amongst cooperating individuals. The mechanism underlying this differentiation in plasticity of cooperative behaviour between the two selection lines used in this experiment is still unclear. Differences in cooperative behaviour are thought to be underpinned by more general traits, such as responsiveness to social stimuli (16). While previous work using the same phenotypic selection lines has found no differences in overall sociability (time spent with conspecifics), there is some evidence that, at least in males, some aspects of social behaviour, such as social selectivity (rate of social sampling), differ (27). More research is needed to understand the full extent of divergence in behaviour between individuals of differing cooperative phenotype, and the effects such differences may have on the motivation for and the flexibility of cooperative behaviour.

Our model demonstrated that more cooperative individuals can evolve to show higher plasticity compared to less cooperative individuals, even under very simple behavioural and evolutionary rules. This suggests that the behaviour that we observed in this experiment can arise via simple evolutionary processes, and does not rely on high levels of cognition that other mechanisms of assortment might rely on (e.g., reputational book-keeping (35)). We also found that the relative cost incurred when nobody cooperates affected the strength of the relationship between cooperation and plasticity, suggesting that ecological factors could be important in the relationship between these traits. Guppies present a unique opportunity to study these relationships because they naturally occur under different levels of predation threat, which represents a key driver of guppies’ cooperative behaviour (36). Empirical studies of wild guppies from different predation regimes could help to uncover the importance of the link between cooperation, plasticity and the ecological drivers of cooperative behaviour in the wild.

Interestingly, while we found that more cooperative (but not less cooperative) fish modified the time spent inspecting the predator, i.e. the amount of inspection, as a function of their past and current social partner’s behaviour, we found that average distance of inspection, i.e. its intensity, did not change across consecutive trials in either selection line. A collapsed measure of amount and intensity of inspection in mirrored trials has been shown to positively correlate with time spent occupying the lead position during inspection with a live partner in wild-caught fish (8). This supports our use of these measures as representative of cooperative investment, but suggests that these two elements of investment should be considered separately. The biological significance of the differentiation between timing and distance of inspection is unclear without further work and we must keep in mind that it is possible that we failed to detect plasticity in intensity as there was generally greater variation in this measure. However, it may be the case that time spent inspecting reflects greater cost moderation compared to average distance of inspection, and therefore is a more important proxy for cooperative investment than the distance to the predator stimulus. It is also possible that the two selection lines do not differ in how close they inspect the predator stimulus when their partner matches them in investment (i.e. the mirror image will never be able to over- or under-take the focal individual) leading to high coordination. When these two selection lines were generated, the distance of inspection from the predator differed between more cooperative (HL) and less cooperative (LL) fish (27). The standardizing effect on partner coordination (i.e. the distance between them during an inspection) (37) with the use of a mirror in the current experiment instead of Robofish (which was used in 27) as a simulated social partner is likely to be an important underlying factor explaining this difference between the studies. Given that leading during inspection is more costly than lagging (23), coordination and adjustment of relative position is likely to be an important factor affecting the cost of predator inspection, and this could affect how closely individuals inspect, and the extent to which they adjust this. This is supported by the finding that guppies originating from locales under severe predation pressure exhibit greater coordination between inspecting fish compared to fish originating from low predation habitats (37).

Fish from both more cooperative and less cooperative lines adjusted their predator inspection behaviour as a function of the behaviour of their social partners; this is in agreement with past research by Edenbrow and colleagues (37), who found that current and previous cooperative experiences affect an individual’s own cooperative behaviour. We also confirmed Edenbrow et al.’s finding that the current social partner’s behaviour has a more pronounced effect on cooperative behaviour compared to previous social partners, although only in individuals from the more cooperative line and only when current partners cooperated following a prevailing experience of defection from social partners. Our findings suggest that, while cooperative behaviour is affected by past social experiences and, crucially, the cooperative behaviour of the current social partner, the extent to which an individual reciprocates the behaviour of a partner depends on its own behavioural traits.

We observed no effect of sex on the plasticity of cooperative behaviour in either phenotypic selection line (see Supplementary Information). This may appear counterintuitive, given both the documented sex differences in predator inspection behaviour in this species (e.g. 38) and the differential risk experienced by inspecting female and male fish (39,40), as well as the documented differences in aggressiveness and social selectivity between males originating from the two phenotypic selection lines used in this study (27). In this study, we simulated a novel social partner during each predator exposure, rather than familiar partners (although it cannot be precluded that the focal individual may form some kind of context-dependent familiarity with their simulated social partner – see (41)). This represents only one of the social contexts that particularly female guppies will cooperate under given the documented social structure in natural populations that might otherwise influence sex differences in inspection behaviour. Female guppies – but not males – form stable and long-lasting associations (42) where consistency in cooperative behaviour would appear beneficial to net fitness (16), and females associate preferentially with familiar females (43). It is, therefore, possible that males and females do not differ in their investment decisions in contexts that do not involve an established familiarity, such as inspecting with a mirror image (as also shown in 26). Further experimental work is needed to determine if, given the opportunity for repeated interaction with the same social partners, plasticity in cooperative behaviour would differ between male and female guppies.

Our experimental results demonstrate that more cooperative individuals are more responsive to their immediate social environment than less cooperative individuals in the context of predator inspection in wild-type Trinidadian guppies. Our model highlights that the coevolution of cooperation and plasticity in cooperative behaviour can arise by very simple behavioural rules, and is likely affected by ecological factors. Taken together, these findings suggest a mechanism by which positive assortment amongst cooperative individuals can be directly supported and that could potentially augment other mechanisms driving assortment (e.g., walk away (44)), and how this might be impacted by ecological factors, such as predation threat. Further work is now required to identify systems in which coevolution between cooperative phenotype and plasticity can facilitate the evolution of cooperation.

## Acknowledgements

The authors would like to thank Christine Soper, Adam Johnstone and Jake Cranston for help with animal husbandry, as well as Bridie Barton and Samuel James for help with data collection.

## Authors’ contributions

**Sylvia Dimitriadou:** Conceptualisation, Methodology, Investigation, Writing – original draft. **Rebecca Padget:** Methodology, Investigation, Formal Analysis, Writing – original draft, Visualisation. **Tegen Jack:** Investigation, Writing – review & editing. **Safi Darden:** Conceptualisation, Methodology, Investigation, Supervision, Funding acquisition, Writing – review & editing.

## Data and code availability

The data and code for this manuscript will be made available on GitHub upon acceptance for publication.

## References

1. Bshary R, Oliveira RF. Cooperation in animals: Toward a game theory within the framework of social competence. Curr Opin Behav Sci. 2015;3:31–7.

2. Clutton-Brock T. Cooperation between non-kin in animal societies. Nature. 2009;462(7269):51–7.

3. Taborsky M, Frommen JG, Riehl C. Correlated pay-offs are key to cooperation. Philos Trans R Soc Lond B Biol Sci. 2016 Feb 5;371(1687):20150084.

4. Lehmann L, Keller L. The evolution of cooperation and altruism – a general framework and a classification of models. J Evol Biol. 2006 Sep 1;19(5):1365–76.

5. Nowak MA. Evolving cooperation. J Theor Biol. 2012;299(0):1–8.

6. Sachs JL, Mueller UG, Wilcox TP, Bull JJ. The evolution of cooperation. Q Rev Biol. 2004;79(2):135–60.

7. Fletcher JA, Doebeli M. A simple and general explanation for the evolution of altruism. Proc Biol Sci. 2009 Jan 7;276(1654):13–9.

8. Brask JB, Croft DP, Edenbrow M, James R, Bleakley BH, Ramnarine IW, et al. Evolution of non-kin cooperation: social assortment by cooperative phenotype in guppies. R Soc Open Sci. 2019 Jan 31;6(1):181493.

9. Trivers RL. The evolution of reciprocal altruism. Q Rev Biol. 1971;46(1):35–57.

10. Hamilton WD, Axelrod R. The evolution of cooperation. Science. 1981;211(27):1390–6.

11. van--Doorn GS, Taborsky M. The evolution of generalized reiprocity on social interaction networks. Evolution. 2011; Early View: doi:10.1111/j.1558-5646.2011.01479.x.

12. Eshel I, Cavalli-Sforza LL. Assortment of encounters and evolution of cooperativeness. Proc Natl Acad Sci. 1982;79(4):1331–5.

13. Dakin R, Ryder TB. Dynamic network partnerships and social contagion drive cooperation. Proc R Soc B [Internet]. 2018 Dec 19 [cited 2022 Jul 27];285(1893). Available from: https://royalsocietypublishing.org/doi/10.1098/rspb.2018.1973

14. Dingemanse NJ, Kazem AJN, Réale D, Wright J. Behavioural reaction norms: animal personality meets individual plasticity. Trends Ecol Evol. 2010 Feb 1;25(2):81–9.

15. Nowak MA. Five rules for the evolution of cooperation. Science. 2006 Dec 8;314(5805):1560–3.

16. Bergmüller R, Schürch R, Hamilton IM. Evolutionary causes and consequences of consistent individual variation in cooperative behaviour. Philos Trans R Soc Lond B Biol Sci. 2010 Sep 12;365(1553):2751–64.

17. Rand DG, Arbesman S, Christakis NA. Dynamic social networks promote cooperation in experiments with humans. Proc Natl Acad Sci. 2011 Nov 29;108(48):19193–8.

18. Van--Segbroeck S, Santos FC, Nowé A, Pacheco JM, Lenaerts T. The evolution of prompt reaction to adverse ties. BMC Evol Biol. 2008;8(1):1–8.

19. Allan JR, Pitcher TJ. Species segregation during predator evasion in cyprinid fish shoals. Freshw Biol. 1986;16(5):653–9.

20. Magurran AE, Seghers BH. Predator inspection behaviour covaries with schooling tendency amongst wild guppy, Poecilia reticulata, populations in Trinidad. Behaviour. 1994;128(1):121–34.

21. Pitcher TJ, Green DA, Magurran AE. Dicing with death: predator inspection behaviour in minnow shoals. J Fish Biol. 1986 Apr 1;28(4):439–48.

22. Milinski M. Tit for tat in sticklebacks and the evolution of cooperation. nature. 1987;325(6103):433–5.

23. Milinski M, Lüthi JH, Eggler R, Parker GA. Cooperation under predation risk: experiments on costs and benefits. Proc R Soc Lond B Biol Sci. 1997;264(1383):831–7.

24. Pitcher TJ. Who dares, wins: the function and evolution of predator inspection behaviour in shoaling fish. Neth J Zool. 1991;42(2):371–91.

25. Dugatkin LA. Tendency to inspect predators predicts mortality risk in the guppy (Poecilia reticulata). 1992;3(2):124–7.

26. Dugatkin LA. Do guppies play TIT FOR TAT during predator inspection visits? Behav Ecol Sociobiol. 1988;23(6):395–9.

27. Dimitriadou S, Croft DP, Darden SK. Divergence in social traits in Trinidadian guppies selectively bred for high and low leadership in a cooperative context. Sci Rep. 2019;9(1):1–12.

28. Croft DP, Arrowsmith BJ, Bielby J, Skinner K, White E, Couzin ID, et al. Mechanisms underlying shoal composition in the Trinidadian guppy, Poecilia reticulata. Oikos. 2003;100(3):429–38.

29. De Santi A, Sovrano VA, Bisazza A, Vallortigara G. Mosquitofish display differential left-and right-eye use during mirror image scrutiny and predator inspection responses. Anim Behav. 2001;61(2):305–10.

30. Dugatkin LA, Alfieri M. Guppies and the TIT FOR TAT strategy: pREFerence based on past interaction. Behav Ecol Sociobiol. 1991 Apr;28(4):243–6.

31. Hesse S, Anaya-Rojas JM, Frommen JG, Thünken T. Kinship reinforces cooperative predator inspection in a cichlid fish. J Evol Biol. 2015;28(11):2088–96.

32. Bürkner PC. brms: An R Package for Bayesian Multilevel Models Using Stan. J Stat Softw. 2017 Aug 29;80:1–28.

33. R Development Core Team. R: A Language and Environment for Statistical Computing. Vienna, Austria: R Foundation for Statistical Computing; 2014. R Foundation for Statistical Computing. … Free Available Internet Httpwww R-Proj … [Internet]. 2015; Available from: https://scholar.google.co.in/scholar?hl=en&q=R+core+team+&btnG=#1%5Cnhttps://scholar.google.co.in/scholar?hl=en&q=R+core+team+&btnG=%238%5Cnhttps://scholar.google.co.in/scholar?hl=en&q=R+core+team+&btnG=%232

34. Arel-Bundock V, Diniz MA, Greifer N. Marginaleffects: Marginal Effects, Marginal Means, Predictions, and Contrasts. R Package Version 05 0 URL Httpsvincentarelbundock Github Iomarginaleffects. 2022;

35. Melis AP, Raihani NJ. The cognitive challenges of cooperation in human and nonhuman animals. Nat Rev Psychol. 2023 Sep;2(9):523–36.

36. Magurran AE. Evolutionary ecology: the Trinidadian guppy. Vol. 19. Oxford University Press on Demand; 2005. 224 p.

37. Edenbrow M, Bleakley BH, Darden SK, Tyler CR, Ramnarine IW, Croft DP. The Evolution of Cooperation: Interacting Phenotypes among Social Partners. Am Nat. 2017;189(6):000–000.

38. Magurran AE, Nowak MA. Another Battle of the Sexes: The Consequences of Sexual Asymmetry in Mating Costs and Predation Risk in the Guppy, Poecilia reticulata. Proc R Soc B Biol Sci. 1991;246(1315):31–8.

39. Seghers BH. Analysis of geographic variation in the antipredator adaptations of the guppy : Poecilia reticulata. 1973 [cited 2018 Feb 5]; Available from: https://open.library.ubc.ca/cIRcle/collections/ubctheses/831/items/1.0100947

40. Rodd FH, Reznick DN. Variation in the demography of guppy populations: the importance of predation and life histories. Ecology. 1997;78(2):405–18.

41. Griffiths SW, Magurran AE. Schooling decisions in guppies ( Poecilia reticula are based matching by phenotype on familiarit rather than kin recognition. Behav Ecol Sociobiol. 1999;45(6):437–43.

42. Croft DP, Morrell LJ, Wade AS, Piyapong C, Ioannou CC, Dyer JRG, et al. Predation risk as a driving force for sexual segregation: a cross-population comparison. Am Nat. 2006 Jun 17;167(6):867–78.

43. Griffiths SW, Magurran AE. Sex and schooling behaviour in the Trinidadian guppy. Anim Behav. 1998;56(3):689–93.

44. Aktipis CA. Know when to walk away: Contingent movement and the evolution of cooperation. J Theor Biol. 2004;231(2):249–60.

