## Supplementary Information for "Going with the flow: evidence for phenotypic variation in cooperative plasticity during predator inspection in Trinidadian guppies (*Poecilia reticulata*)"

### Supplemental Information

#### Methods

#### Behavioural assay

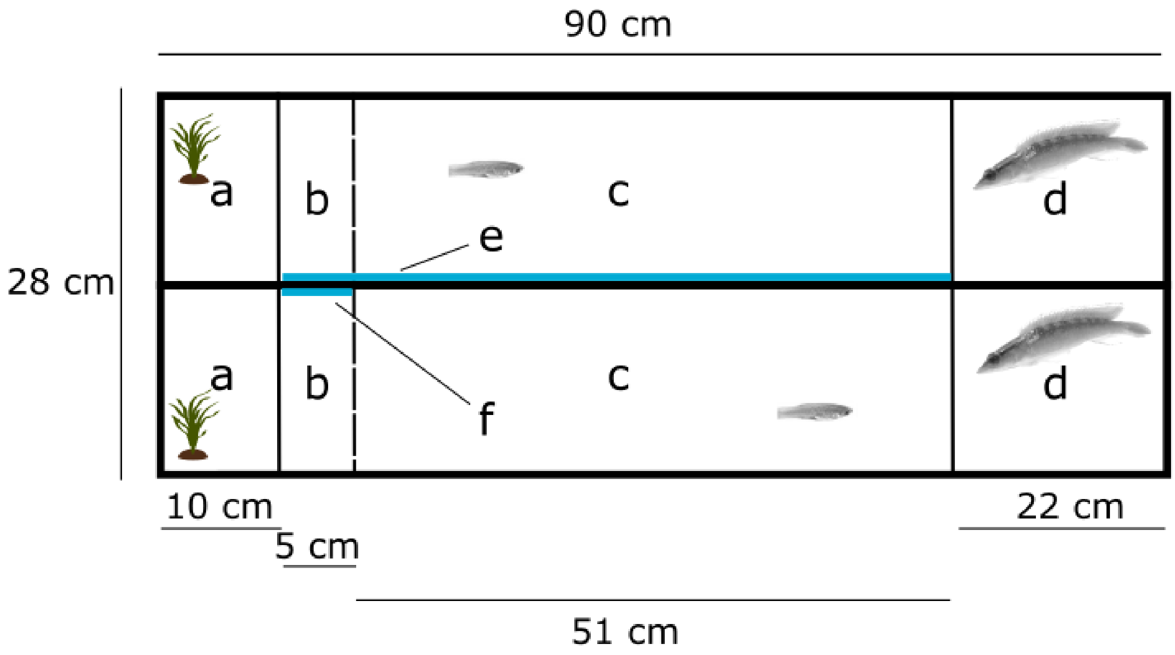


**Figure S1. Experimental setup for the behavioural assay (top view).** The customised arena comprised two inspection lanes separated by a watertight opaque barrier. a: plant compartment; b: refuge area; c: inspection lane; d: predator compartment; e: full-length mirror simulating cooperation; f: short mirror simulating defection.

**Table S1. Experimental treatments and sample sizes.** Four individuals were removed from the analysis (see text).

| Selection line | Experimental Phase | Test Phase | Female (n) | Male (n) |
|---|---|---|---|---|
| High Leading | Cooperation | Cooperation | 11 | 10 |
| High Leading | Cooperation | Defection | 10 | 11 |
| High Leading | Defection | Cooperation | 10 | 10 |
| High Leading | Defection | Defection | 10 | 10 |
| Low Leading | Cooperation | Cooperation | 9 | 8 |
| Low Leading | Cooperation | Defection | 10 | 11 |
| Low Leading | Defection | Cooperation | 10 | 10 |
| Low Leading | Defection | Defection | 10 | 10 |

**Results: main results**

**Table S2. Effect sizes (marginal effect) and credible intervals for contrasts of interest for time spent inspecting (groups compared are shown in bold text)**. Both marginal effects and confidence intervals are given in seconds. Where credible intervals do not overlap zero (indicating evidence of a possible effect), we show results in italics.

| Comparison | Marginal effect | Credible interval |
|---|---|---|
| High-leadership under unexpected cooperation compared to exposure-phase defection (HL: DDD/**C** v **DDD**/C) | +39 seconds | *50% = 17, 63*<br>89% = -12, 93<br>95% = -24, 102 |
| Low-leadership under unexpected cooperation compared to exposure-phase defection (LL: DDD/**C** v **DDD**/C) | -2.4 seconds | 50% = -24, 21<br>89% = -54, 51<br>95% = -63, 63 |
| High-leadership under unexpected defection compared to exposure-phase cooperation (HL: CCC/**D** v **CCC**/D) | -29 seconds | 50% = -51, 6<br>89% = -81, 24<br>95% = -90, 36 |
| Low-leadership under unexpected defection compared to exposure-phase cooperation (LL: CCC/**D** v **CCC**/D) | -13 seconds | 50% = -36, 9<br>89% = -66, 36<br>95% = -81, 48 |

| | | |
|---|---|---|
| High-leadership under unexpected compared to expected cooperation (HL: DDD/**C** v **CCC/C**) | +30 seconds | *50% = 18, 42*<br>89% = -1.1, 63<br>95% = -8.7, 69 |
| Low-leadership under unexpected compared to expected cooperation (LL: DDD/**C** v **CCC/C**) | -36 seconds | *50% = –51, -23*<br>*89% = –72, -3*<br>95% = –81, 4.5 |
| High-leadership under unexpected compared to expected defection (HL: CCC/**D** v **DDD/D**) | -33 seconds | *50% = -48, -18*<br>89% = -69, 2.7<br>95% = -75, 11 |
| Low-leadership under unexpected compared to expected defection (LL: CCC/**D** v **DDD/D**) | -7 seconds | 50% = -23, 8.7<br>89% = -45, 30<br>95% = -54, 36 |

**Table S3. Effect sizes (marginal effect) and credible intervals for contrasts of interest for distance to the predator (groups compared are shown in bold text)**. Both marginal effects and confidence intervals are given on the scale of the data. Where credible intervals do not overlap zero (indicating evidence of a possible effect), we show results in italics.

| **Comparison** | **Marginal effect** | **Credible interval** |
|---|---|---|
| High-leadership under unexpected cooperation compared to exposure-phase defection (HL: DDD/**C** v **DDD**/C) | +1.7cm | 50% = -15, 23<br>89% = -41, 64<br>95% = -50, 84 |
| Low-leadership under unexpected cooperation compared to exposure-phase defection (LL: DDD/**C** v **DDD**/C) | -4.8 cm | 50% = -30, 7.8<br>89% = -70, 33<br>95% = -89, 43 |
| High-leadership under unexpected defection compared to exposure-phase cooperation (HL: CCC/**D** v **CCC**/D) | +6.1 cm | 50% = -18, 4<br>89% = -44, 18<br>95% = -57, 24 |
| Low-leadership under unexpected defection compared to exposure-phase cooperation (HL: CCC/**D** v **CCC**/D) | +5.0 cm | 50% = -14, 4<br>89% = -26, 19<br>95% = -32, 26 |
| High-leadership under unexpected compared to expected cooperation (HL: DDD/**C** v **CCC/C**) | -4.1 cm | *50% = -14, -3.1*<br>89% = -22, 4.6<br>95% = -25, 8.4 |

| | | |
|---|---|---|
| Low-leadership under unexpected compared to expected cooperation (LL: DDD/**C** v **CCC/C**) | +4.6 cm | *50% = 3.8, 17*<br>89% = -3.2, 30<br>95% = -6.4, 39 |
| High-leadership under unexpected compared to expected defection (HL: CCC/**D** v **DDD/D**) | +6.1 cm | *50% = 5.1, 20*<br>89% = -2.8, 38<br>95% = -6.3, 47 |
| Low-leadership under unexpected compared to expected defection (LL: CCC/**D** v **DDD/D**) | +3.1 cm | *50% = 1.0, 13*<br>89% = -6.2, 24<br>95% = -8.9, 29 |

**Results: other model parameters**

**Table S4. Estimates and 95% credible intervals (CI) for the other parameters in the time inspecting and distance from predator models.**

| **Model** | **Parameter** | **Estimate** | **95% CI** |
|---|---|---|---|
| Time inspecting | Sex | 0.26 | 0.03, 0.50 |
| | Lane 2a | -0.26 | -0.77, 0.24 |
| | Lane 3a | 0.18 | -0.25, 0.61 |
| | Lane 4a | -0.20 | -0.72, 0.32 |
| | Lane 1b | 0.20 | -0.20, 0.62 |
| | Lane 2b | -0.25 | -0.73, 0.23 |
| | Lane 3b | 0.21 | -0.24, 0.66 |
| | Lane 4b | -0.31 | -0.80, 0.19 |
| | Predator | 0.06 | -0.12, 0.24 |
| | Trial day (2) | -0.26 | -0.50, -0.03 |
| | Trial day (3) | -0.13 | -0.37, 0.13 |
| | Trial day (4) | -0.09 | -0.40, 0.21 |
| Distance from predator | Sex | -0.07 | -0.15, 0.00 |
| | Lane 2a | 0.04 | -0.14, 0.21 |
| | Lane 3a | 0.00 | -0.15, 0.14 |
| | Lane 4a | 0.08 | -0.11, 0.28 |
| | Lane 1b | -0.03 | -0.17, 0.11 |
| | Lane 2b | -0.01 | -0.18, 0.16 |
| | Lane 3b | 0.00 | -0.15, 0.15 |
| | Lane 4b | 0.10 | -0.07, 0.28 |
| | Predator | -0.02 | -0.09, 0.05 |
| | Trial day (2) | -0.04 | -0.13, 0.05 |
| | Trial day (3) | 0.11 | 0.01, 0.20 |

| | | | |
|---|---|---|---|
| | Trial day (4) | -0.10 | -0.23, 0.03 |

**Post-hoc test for sex differences in plasticity**

Because our model identified evidence that males inspected closer and for longer than females, we ran an additional analysis to identify whether there might be differences in plasticity between males and females. To do this, we included an additional term for the interaction between sex, social environment and condition in the model.

We found no evidence that sexes differed in their plasticity for either time spent inspecting or distance of inspections (Table S2).

**Table S5. Effect sizes (marginal effect) and credible intervals for contrasts of interest for time spent inspecting (groups compared are shown in bold text). Both marginal effects and confidence intervals are given on the scale of the data. Where credible intervals do not overlap zero (indicating evidence of a possible effect), we show results in italics.**

| | **Comparison** | **Marginal effect** | **Credible intervals** |
|---|---|---|---|
| Time spent inspecting | Females under unexpected vs expected cooperation (F: DDD/**C** v **CCC/C**) | 15 seconds | 50% = -6, 33<br>89% = -36, 57<br>95% = -48, 69 |
| | Males under unexpected vs expected cooperation (M: DDD/**C** v **CCC/C**) | -15 seconds | 50% = -36, 6<br>89% = -63, 33<br>95% = -60, 45 |
| | Females under unexpected vs expected defection (F: CCC/**D** v **DDD/D**) | -9 seconds | 50% = -30, 12<br>89% = -60, 42<br>95% = -72, 54 |
| | Males under unexpected vs expected defection (M: CCC/**D** v **DDD/D**) | -18 seconds | 50% = -39, 6<br>89% = -69, 36<br>95% = -81, 45 |
| | Females under unexpected cooperation vs exposure-phase defection (F: DDD/**C** v **DDD**/C) | -0.9 seconds | 50% = -15, 15<br>89% = -90, 36<br>95% = -48, 42 |
| | Males under unexpected cooperation vs exposure-phase defection (M: DDD/**C** v **DDD**/C) | 6 seconds | 50% = -9, 21<br>89% = -33, 39<br>95% = -42, 51 |
| | Females under unexpected defection vs exposure-phase cooperation (F: CCC/**D** v **CCC**/D) | -21 seconds | 50% = -45, 1<br>89% = -78, 33<br>95% = -87, 45 |
| | Males under unexpected defection vs exposure-phase cooperation (M: CCC/**D** v **CCC**/D) | -33 seconds | *50% = -54, -9*<br>89% = -87, 24<br>95% = -96, 36 |
| Distance of inspections | Females under unexpected vs expected cooperation (F: DDD/**C** v **CCC/C**) | -6 cm | 50% = -9, 2<br>89% = -13, 2<br>95% = -15, 4 |

| | | | |
|---|---|---|---|
| | Males under unexpected vs expected cooperation (M: DDD/**C** v **CCC/C**) | -2 cm | 50% = -4, 3<br>89% = -8, 8<br>95% = -10, 10 |
| | Females under unexpected vs expected defection (F: CCC/**D** v **DDD/D**) | -2 cm | 50% = -5, 2<br>89% = -9, 7<br>95% = -11, 9 |
| | Males under unexpected vs expected defection (M: CCC/**D** v **DDD/D**) | -0.3 cm | 50% = -3, 4<br>89% = -8, 13<br>95% = -10, 10 |
| | Females under unexpected cooperation vs exposure-phase defection (F: DDD/**C** v **DDD**/C) | -5 cm | 50% = -8, 3<br>89% = -11, 5<br>95% = -12, 2 |
| | Males under unexpected cooperation vs exposure-phase defection (M: DDD/**C** v **DDD**/C) | -2 cm | 50% = -5, 0.2<br>89% = -8, 4<br>95% = -9, 5 |
| | Females under unexpected defection vs exposure-phase cooperation (F: CCC/**D** v **CCC**/D) | 5 cm | *50% = 1, 9*<br>89% = -4, 14<br>95% = -6, 16 |
| | Males under unexpected defection vs exposure-phase cooperation (M: CCC/**D** v **CCC**/D) | 5 cm | *50% = 2, 9*<br>89% = -4, 14<br>95% = -6, 16 |
